# Gut microbiota diversity and C-Reactive Protein are predictors of disease severity in COVID-19 patients

**DOI:** 10.1101/2021.04.20.440658

**Authors:** André Moreira-Rosário, Cláudia Marques, Hélder Pinheiro, João Ricardo Araújo, Pedro Ribeiro, Rita Rocha, Inês Mota, Diogo Pestana, Rita Ribeiro, Ana Pereira, Maria José de Sousa, José Pereira-Leal, José de Sousa, Juliana Morais, Diana Teixeira, Júlio César Rocha, Marta Silvestre, Nuno Príncipe, Nuno Gatta, José Amado, Lurdes Santos, Fernando Maltez, Ana Boquinhas, Germano de Sousa, Nuno Germano, Gonçalo Sarmento, Cristina Granja, Pedro Póvoa, Ana Faria, Conceição Calhau

**Author notes:** Corresponding author Conceição Calhau, Faculdade de Ciências Médicas|NOVA Medical School, Universidade Nova de Lisboa, Campo Mártires da Pátria, 130, 1169-056 Lisboa, Portugal. These authors contributed equally to this study (André Moreira-Rosário, Cláudia Marques).

## Abstract

Risk factors for COVID-19 disease severity are still poorly understood. Considering the pivotal role of gut microbiota on host immune and inflammatory functions, we investigated the association between changes in gut microbiota composition and the clinical severity of COVID-19. We conducted a multicentre cross-sectional study prospectively enrolling 115 COVID-19 patients categorized according to: 1) WHO Clinical Progression Scale - mild 19 (16.5%), moderate 37 (32.2%) or severe 59 (51.3%); and 2) location of recovery from COVID-19 - ambulatory 14 (household isolation; 12.2%), hospitalized in ward 40 (34.8%) or intensive care unit 61 (53.0%). Gut microbiota analysis was performed through 16S rRNA gene sequencing and data obtained was further related with clinical parameters of COVID-19 patients. Risk factors for COVID-19 severity were identified by univariate and multivariable logistic regression models.

In comparison with mild COVID-19 patients, the gut microbiota of moderate and severe patients has: a) lower Firmicutes/Bacteroidetes ratio, b) higher abundance of Proteobacteria; and c) lower abundance of beneficial butyrate-producing bacteria such as *Roseburia* and *Lachnospira* genera. Multivariable regression analysis showed that Shannon index diversity (odds ratio [OR] 2.85 [95% CI 1.09-7.41]; p=0.032) and C-Reactive Protein (OR 3.45 [95% CI 1.33-8.91]; p=0.011) were risk factors for COVID-19 severe disease (a score of 6 or higher in WHO clinical progression scale).

In conclusion, our results demonstrated that hospitalised moderate and severe COVID-19 patients have microbial signatures of gut dysbiosis and for the first time, the gut microbiota diversity is pointed out as a prognostic biomarker for COVID-19 disease severity.

## Introduction

COVID-19 caused by the novel coronavirus SARS-CoV-2 infection, is clinically diverse in terms of disease severity – ranging from absence of symptoms, to mild, self-limiting respiratory illness (including the common cold), severe pneumonia, acute respiratory distress syndrome and death (1). COVID-19-induced respiratory distress syndrome was described to be associated with exuberant inflammation, intense cytokine production (cytokine storm syndrome) and multi-organ dysfunction (1, 2). Although respiratory symptoms are the most commonly reported among COVID-19 patients, gastrointestinal symptoms are also likely between SARS-CoV-2 infected patients indicating that the gastrointestinal tract is as well an infected organ (3). In consequence, SARS-CoV-2 is detected in faeces of some COVID-19 patients (4-6).

Although risk groups for severe COVID-19 disease were identified as being primarily the elderly and individuals with comorbidities, such as hypertension and diabetes (7-9), COVID-19 may evolve adversely even in individuals without comorbidities, causing severe pneumonia, long-term sequelae and eventually death(10). These observations suggest the existence of major predisposition factor(s) related with disease progression that need(s) to be urgently unveiled.

The human gut microbiota mainly composed by bacteria, plays a critical role in health and most notably in host immune response, including vaccine efficacy (11, 12). Changes in gut microbiota composition have been reported to affect both vulnerability and disease outcomes in non-communicable diseases, such as diabetes, inflammatory bowel disease, and obesity, leading to a state of chronic low-grade inflammation (13-15). This role of gut microbiota in both immune and inflammatory responses, together with the fact that SARS-CoV-2 binds to angiotensin converting enzyme (ACE) 2 receptors on gut epithelium (16) where it has been detected along the faeces of COVID-19 patients (17), suggest the existence of a microbial fingerprinting among these patients that may provide a predictive value for disease severity. Accordingly, gut microbiome characterization has been assessed in COVID-19 patients that unveiled profound alterations on bacterial composition (4, 18, 19). The depletion of beneficial bacteria from *Lachnospiraceae* taxa and *Bifidobacterium, Faecalibacterium* and *Roseburia* genera (18-20) has been proposed as having an impact on the modulation of host immune response to SARS-CoV-2 infection and potentially influenced disease severity and outcomes (18). However, existing studies did not enrol COVID-19 patients representative of the different COVID-19 severity levels, lacking mainly patients with severe clinical manifestations. Most importantly, previous studies did not clarify whether the observed changes in microbiota composition are a common patient’s response to SARS-CoV-2 infection rather than directly involved in disease severity.

Taking this into consideration, we investigate the association between gut microbiota and COVID-19 disease severity using a cohort of 115 patients stratified by asymptomatic/mild-moderate-severe according with the WHO Clinical Progression Scale. Considering that previous studies have shown that alterations in gut microbiota do not alter significantly during COVID-19 disease progression and even after SARS-CoV-2 clearance (18, 19), one point faecal collection was performed and clinical variables and gut bacterial composition were compared between COVID-19 severity groups. The role of antibiotic use was also addressed. To the best of our knowledge, this is the largest study to assess the gut microbiota composition in patients with COVID-19, and the first outside of China.

## Materials and methods

### Study design and population

This national multicentre cross-sectional study was conducted in six geographically different Portuguese centres selected by invitation. The distribution of patients per participating centre was 38 (33.0%), 33 (28.7%), 18 (15.7%), 12 (10.4%), 8 (7.0%), and 6 (5.2%). Patients eligibility criteria included age equal or above 18 years old and a positive test for SARS-CoV-2 by nasopharyngeal swabs using quantitative RT-PCR performed in national reference laboratories and in accordance with recommendations from the National Directorate of Health. COVID-19 patients were recruited during the first wave of pandemic in Portugal - from 21^st^ April 2020 to 1^st^ July 2020 - and sample size was determined based on the feasibility of recruitment during this period. The minimally detectable effect sizes were calculated retrospectively. In order to achieve a statistical power of 80% and a two-sided significance level of 0.05, and considering the total sample size of 115 individuals, the study was powered to detect a mean difference of 0.15 in the Shannon’s Diversity Index between mild-to-moderate and severe COVID-19 patients.

Participating centres prospectively collected data from consecutive patients included in the study and classified them according to location of recovery (ambulatory, hospitalization in ward or intensive care unit [ICU]) and disease severity using the WHO Clinical Progression Scale (21) (mild, moderate and severe). Ethic committees and institutional review boards from participating centres approved the study protocol considering it a minimal-risk research using data collected for routine clinical practice and waived the requirement to obtain informed consent. Patients (or their proxies) received written information about the study and were informed about their right to refuse to participate. The study was registered at ClinicalTrials.gov, number NCT04355741. All authors had access to the study data and reviewed and approved the final version of the manuscript.

### Data collection

Patient demographic characteristics, severity scores, smoking habits, comorbidities prior to hospitalisation (diabetes, hypertension, chronic respiratory diseases, immunosuppression, haematological oncological disease, previous chronic therapy, and others), or antibiotic exposure six months prior to COVID-19 diagnosis were recorded for all patients at baseline (*i*.*e*. immediately after subject enrolment). Data on clinical presentation of COVID-19, C-reactive protein (CRP) levels, antibiotic, antiviral and steroid treatments received during the course of disease, as well as nutritional and respiratory support (as per WHO Clinical Progression Scale (21)) were collected. In addition, clinical outcomes such as duration of mechanical ventilation, ICU length of stay, ICU mortality, and 28-day mortality were also collected. Patients were followed up until hospital discharge if that was the case.

### Stool Collection

Faecal samples of COVID-19 patients were collected after subject enrolment (single point collection). Faecal samples were collected with a stool collection kit (EasySampler, ALPCO) containing RNAlater (Sigma-Aldrich). Faecal samples were kept at −80 °C until nucleic acid extraction.

### Gut microbiota

Genomic DNA was extracted and purified from stool samples of COVID-19 patients using the NZY Tissue gDNA Isolation Kit (NZYTech). All 16S DNA libraries (V3 and V4 regions) were prepared, sequenced and analysed in accordance with the manufacturer’s instructions for each kit and instrument. Briefly, 16S DNA libraries were prepared using the Ion 16S™ Metagenomics Kit targeted panel (Thermo Fisher Scientific) and each sample was individually identified with the Ion Xpress™ Barcode Adapters Kits (Thermo Fisher Scientific). All available regions were amplified using the Ion 16S™ Metagenomics Kit (Thermo Fisher Scientific). Amplified fragments were then prepared for sequencing using the Ion CHEF system (Thermo Fisher Scientific) and loaded into Ion 318 Chip Kit v2 BC (Thermo Fisher Scientific). Sequencing runs were performed on an Ion S5 System (Thermo Fisher Scientific) aiming for a mean sequencing depth coverage of 12000×. Sequencing depths were not normalized in order to achieve a better identification of alpha diversity in each sample. Sequencing data was filtered for length (cutadapt -m 80) and for quality (fastx_trimmer -l 280) after which the V3 and V4 regions were extracted (Mothur align.seqs and screen.seqs). The resulting fastq file was used for taxonomy. The taxonomy of each sample was determined using Kraken2 (https://ccb.jhu.edu/software/kraken2/) and Bracken (https://ccb.jhu.edu/software/bracken) softwares, using our custom 16S database (GutHealth_DB). This database was manually curated by enriching GreenGenes (versions 13_5 and 13_8) with clinically relevant taxa from NCBI RefSeq 16s rRNA sequences (04/2019). The GutHealth_DB currently holds 4765 16s rRNA sequences mapping 1822 species, 1685 genus, 515 families, 404 orders, 248 classes and 89 phyla, and is available upon request. Bacterial species were identified as pathogens or commensals according to The National Microbial Pathogen Database Resource (NMPDR) (https://www.patricbrc.org/view/Taxonomy/561#view_tab=genomes).

### Detection of SARS-CoV-2 in faeces

The following steps were taken to detect SARS-CoV-2 in faeces: 1) RNA extraction by the NucliSENS easyMAG technology based on the Boom technique that utilizes magnetic silica particles from 200-300 mg of stool, and b) detection of SARS-CoV-2 extracted RNA by the EURORealTime SARS-CoV-2 test. The latter is based on reverse transcription to convert viral RNA into complementary DNA, followed by PCR amplification and fluorescence-based real-time detection of two defined sections within the ORF1ab- and N-genes of the SARS-CoV-2 genome. Reverse transcription, amplification and detection of SARS-CoV-2 cDNA were carried out by means of SARS-CoV-2-specific primers and probes.

### Statistical analysis

Statistical analysis was performed using the SPSS version 27 software (SPSS Inc.) and R statistical software package, version V.3.5.1. Descriptive statistics are presented as numbers and percentages for categorical variables, as the mean and standard deviation (SD) for continuous variables or as the medians with interquartile ranges (IQRs) if the continuous variable is not normally distributed. Parametric tests (Student’s t test and one factor analysis of variance-ANOVA) and nonparametric tests (Mann-Whitney and Kruskal-Wallis tests) were used as appropriate, taking into account normality assumptions and the number of groups compared. The Kolmogorov-Smirnov test was used to test normality assumptions of the variable distributions. Chi-square test and Fisher’s exact test were used as appropriate, for categorical variables.

Univariate and multivariate weighted logistic regression models were used in order to evaluate risk factors associated with the severity of COVID-19 (a score of 6 or more in WHO Clinical Progression Scale). The dependent variable in all models was the severity of COVID-19. Independent variables are indicated in table legends (**Table 2**). The Hosmer-Lemeshow statistic and test was applied to evaluate the goodness-of-fit. The discriminative/predictive power of the model was evaluated by the ROC-receiver operating characteristic-curve analysis. The influence of outlier data values on model fit was estimated using leverage statistics, and collinearity was assessed by evaluation of the coefficients’ correlation matrix. The results are presented as crude and adjusted Odds Ratios (OR) and their respective 95% confidence intervals. The statistical significance level was set at 5% and differences were considered statistically significant when p<0.05.

**Table 2.** Bivariate logistic regression analysis of clinical variables associated with severity of COVID-19 (a score of 6 or more in WHO Clinical Progression Scale)

| Variable |  | Crude<sup>a</sup> OR<sup>b</sup> (95% CI) | *p* Value | Adjusted<sup>a</sup> OR<sup>b</sup> (95% CI) | *p* Value |
| --- | --- | --- | --- | --- | --- |
| **Gender** | Female (*n* = 42) | 1.0 | **0.033** |  |  |
|  | Male (*n* = 73) | 2.33 (1.07-5.07) |  |  |  |
| **Age** | <65 yr (*n* = 48) | 1.0 | 0.603 |  |  |
|  | ≥65 (*n* = 67) | 0.82 (0.39-1.73) |  |  |  |
| **C-Reactive Protein** | <96.8 mg/l (*n* = 58) | 1.0 | **0.022** | 1.0 | **0.011** |
|  | ≥96.8 mg/l (*n* = 41) | 2.73 (1.15-6.46) |  | 3.45 (1.33-8.91) |  |
| **Shannon Diversity Index** | ≥2.25 (*n* = 46) | 1.0 | 0.164 | 1.0 | **0.032** |
|  | <2.25 (*n* = 65) | 1.72 (0.80-3.68) |  | 2.85 (1.09-7.41) |  |
| **Overweight or obese** | BMI < 25 (*n* = 36) | 1.0 | 0.749 |  |  |
|  | BMI ≥ 25 (*n* = 69) | 0.88 (0.39-1.98) |  |  |  |
| **Hypertension** | Normal (*n* = 41) | 1.0 | 0.811 |  |  |
|  | Hypertension (*n* = 67) | 0.91 (0.42-1.99) |  |  |  |
| **Diabetes** | Normal (*n* = 62) | 1.0 | 0.101 |  |  |
|  | Diabetes (*n* = 45) | 1.93 (0.88-4.25) |  |  |  |
| **Antibiotic therapy (last 6 months)** | Without (*n* = 66) | 1.0 | 0.244 |  |  |
|  | With (*n* = 42) | 0.63 (0.29-1.37) |  |  |  |

<sup>a</sup>Crude OR were calculated using univariate weighted logistic regression models. Adjusted OR were calculated using multivariate weighted logistic regression models. Fully adjusted estimates take into account four variables (age, antibiotic therapy at least once in the last 6 months, C-Reactive Protein and Shannon Diversity Index) in the model (*n* = 96); <sup>b</sup>Risk (OR) of severity of COVID-19 (a score of 6 or more in WHO Clinical Progression Scale). 95% CI - 95% confidence interval; OR - odds ratio. BMI, Body Mass Index.

Heat tree visualization of the taxonomic differences between the COVID-19 severity groups was produced using the R package metacoder. Coloring indicates all differences between the median proportion of reads for samples from patients grouped according with the severity of COVID-19 using the WHO Clinical Progression Scale *i*.*e*. mild (score 1-3), moderate (score 4-5), and severe disease (score 6-9), as determined using a Wilcox rank-sum test followed by a Benjamini-Hochberg (FDR) correction for multiple testing.

Alpha diversity was measured by the Shannon’s diversity index that summarizes both the species richness (total number of species) and evenness (abundance distribution across species) within a sample. The distances (or dissimilarity) between samples of the same group were compared to the distances between groups using PERMANOVA test.

### Missing data management

Considering that multiple imputation can give rise to biased results when missing data are not random (22), regression analyses were based on complete data. In addition, a sensitivity analysis was performed using multiple imputation in order to account for missing data, with five imputed datasets and ten iterations. All analysis results were aggregated with Rubin’s rule after appropriate transformation (23).

The sensitivity analysis in which missing clinical variables were imputed by means of model-based multiple imputation, showed similar results to the statistical analysis performed with complete cases (Shannon’s Diversity Index: OR=2.71; 95% CI (1.13– 6.52); p=0.026; CRP: OR=4.42; 95% CI (1.61–12.10); p=0.004).

Since missing data were not equally distributed between hospital datasets, we cannot ignore that missing data are not random. Since missing data at random assumption is not testable, we used complete-case analysis as a better approach because multiple imputation could give rise to biased results. Nevertheless, a sensitivity analysis in which missing outcomes were imputed by multiple imputation were also carried out and this analysis showed similar results, which suggests a limited effect of bias and strengthens the results obtained.

## Results

### Clinical characteristics of COVID-19 patients

A total of 115 adults (median age 68; 63.5% males) with a laboratory confirmed positive test for SARS-CoV-2 were included in our study (**Table 1**). More than half (65.7%) were overweight or obese and, regarding co-morbidities, 45 patients (42.1%) had diabetes, 67 (62.0%) hypertension and 21 (19.6%) chronic respiratory disease (**Table 1**). Concerning antibiotic exposure, 42 patients (38.9%) were administered with antibiotics at least once during the 6 months prior to COVID-19 diagnosis (**Table 1**) and 108 (85.2%) were administered antibiotics during the course of COVID-19. According to the location of recovery, the proportion of patients with diabetes attending the ICU was significantly higher than the proportion of patients with diabetes isolated in ward or in ambulatory (31 *vs* 14 patients, p<0.05). Similarly, the proportion of patients presenting three simultaneous comorbidities (obesity, hypertension, and diabetes) was higher in ICU patients than those isolated in the ward or ambulatory (22 *vs* 5 patients, p<0.05).

**Table 1.** Clinical characteristics of COVID-19 patients

| Characteristic | Total<br>(N = 115) | Mild disease<br>(score 1-3)<br>(N = 19) | Moderate disease<br>(score 4-5)<br>(N = 37) | Severe disease<br>(score 6-9)<br>(N = 59) | *p* Value |
| --- | --- | --- | --- | --- | --- |
| Age, median (IQR) — yr | 68.0 (52.0—76.0) | 61.0 (40.0—73.0) | 71.0 (52.0—79.0) | 66.0 (53.0—76.0) | 0.305<sup>a</sup> |
| Male sex — no. (%) | 73 (63.5) | 6 (31.6) | 24 (64.9) | 43 (72.9) | **0.032<sup>b</sup>** |
| Overweight or obese — no. (%) | 69 (65.7) | 7 (70.0) | 24 (66.7) | 38 (64.4) | 0.749<sup>b</sup> |
| Smoker — no. (%) | 21 (19.8) | 2 (18.2) | 5 (13.9) | 14 (23.7) | 0.467<sup>b</sup> |
| Pneumonia Sars-Cov2 — no. (%) | 84 (83.2) | 2 (25.0) | 24 (70.6) | 58 (98.3) | **<0.001<sup>b</sup>** |
| C-Reactive Protein, median (IQR) — mg/L | 72.0 (28.3—158.9) | 32.2 (17.9—54.5) | 63.5 (11.5—115.6) | 96.8 (34.0—177.0) | 0.063<sup>a</sup> |
| Coexisting conditions — no. (%) |  |  |  |  |  |
| Diabetes | 45 (42.1) | 2 (16.7) | 14 (38.9) | 29 (49.2) | 0.099<sup>b</sup> |
| Hypertension | 67 (62.0) | 4 (33.3) | 27 (73.0) | 36 (61.0) | 0.811<sup>b</sup> |
| Chronic respiratory disease | 21 (19.6) | 2 (16.7) | 5 (13.9) | 14 (23.7) | 0.236<sup>b</sup> |
| Immunosuppression | 11 (10.9) | 1 (12.5) | 4 (11.8) | 6 (10.2) | 0.783<sup>b</sup> |
| Haematological-oncological disease | 9 (8.5) | 2 (16.7) | 3 (8.6) | 4 (6.8) | 0.479<sup>b</sup> |
| Medication history — no. (%) |  |  |  |  |  |
| Previous chronic therapy | 86 (86.9) | 8 (100.0) | 29 (87.9) | 49 (84.5) | 0.403<sup>b</sup> |
| Antibiotic therapy (last 6 months) | 42 (38.9) | 5 (41.7) | 17 (45.9) | 20 (33.9) | 0.243<sup>b</sup> |

Patients were classified in accordance with the WHO Clinical Progression Scale. This scale provides a measure of illness severity across a range from 0 (not infected with SARS-CoV-2) to 10 (dead). Patients were grouped in three categories: mild disease (score 1-3), moderate disease (score 4-5), and severe disease (score 6-9). <sup>a</sup>Kruskal-Wallis test. <sup>b</sup>Chi-square test. IQR, interquartile range.

### Faecal microbiota profile according to COVID-19 severity

From the initial 115 COVID-19 patients, we were able to obtain a sufficient amount of good quality faecal DNA to perform microbial composition based on 16S rRNA gene analysis in 111 patients (96.5%). The gut microbiome of the COVID-19 patients was compared based on the fold-change of relative abundance (medians) for each bacterial genus. For this comparison, the COVID-19 patients were grouped according with the disease severity defined by the WHO Clinical Progression Scale (21). This scale provides a measure of illness severity in which a higher score means higher disease severity. Eighteen COVID-19 patients were classified as asymptomatic/mild (score 1-3); thirty-six were categorized as moderate (4-5) while fifty-seven were severe (score 6-9). Three comparisons were done: **1)** severe versus asymptomatic/mild; **2)** severe versus moderate; **3)** moderate versus asymptomatic/mild (from here referred as mild). In order to determine the relative taxonomic changes at genus-level between COVID-19 severity groups, a heat tree was built for each comparison (**Figure 1A**) in which the terminal nodes correspond to bacterial genera. For the first time, our data shows that differences in gut microbiome occur across all phyla with exception of Synergistetes and Verrucomicrobia, and the relative abundance is in general higher in lesser severe COVID-19 states. The higher number of alterations were observed between mild and moderate COVID-19 patients, and between mild and severe states. Lesser alterations were detected between moderate and severe states of COVID-19. Globally, the relative abundances tend to be higher in mild than in moderate patients; in turn, the relative abundances tend to be higher in moderate than in the severe COVID-19 patients. This decrease tendency from mild-to-moderate-to-severe is observed in the bacterial families Bifidobacteriaceae (*Bifidobacterium* genus) and Coriobacteriaceae (*Collinsella* genus) being statistically significant in Lachnospiraceae family, namely in the *Roseburia* and *Lachnospira* genera (p<0.001, FDR corrected). In the opposite direction, *Ralstonia* genus (Proteobacteria) increases with the COVID-19 severity score (p<0.001, FDR corrected).

**Figure 1.**
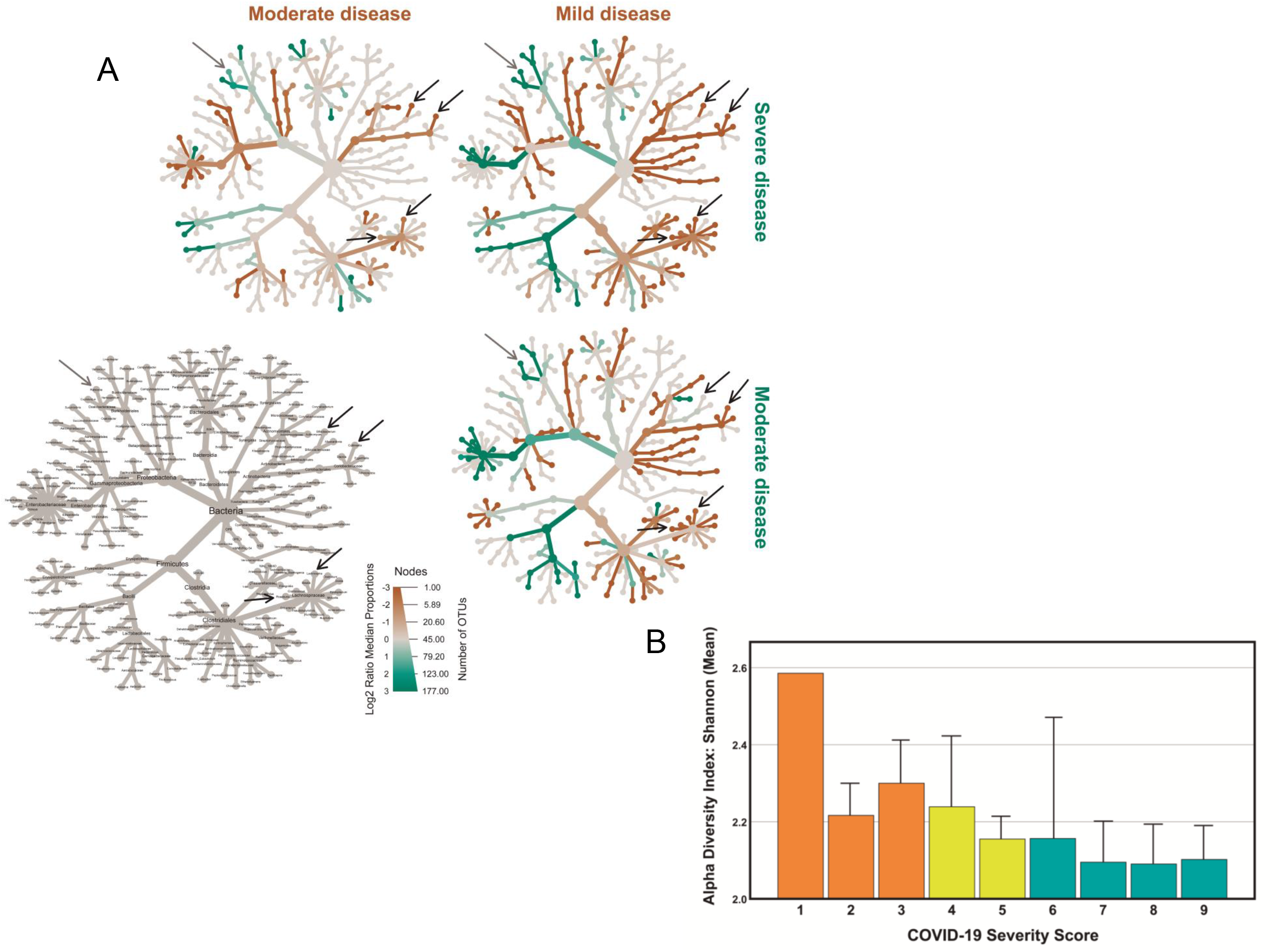
Comparison of COVID-19 gut microbiome with COVID-19 severity. Disease severity was determined according with the WHO Clinical Progression Scale: mild, moderate and severe. **(A)** Heat tree visualization of the taxonomic differences between the COVID-19 severity groups based on Log2 ratio median abundance (proportions), in which the terminal nodes correspond to bacterial genera. The identification of the nodes is shown in the left-bottom image. Three comparisons were done: severe (blue-green) versus mild (orange); severe (blue-green) versus moderate (orange); and ultimately, moderate (blue-green) versus mild (orange). The dominant colour corresponds to a higher number of operational taxonomic units (OTUs); Log2 ratio is 0 (grey colour) when the compared groups are similar. **(B)** Shannon diversity index (mean + SEM) of COVID-19 patients according to WHO Clinical Progression Scale, from score 1 (asymptomatic; viral RNA detected) to score 9 (mechanical ventilation pO2/FiO2 <150 and vasopressors, dialysis, or ECMO).

In accordance with the inverse relation between the relative abundance of bacterial gut microbiota and the COVID-19 severity score, the Shannon’s diversity index shows a similar tendency being higher in mild COVID-19 patients than in moderate and severe, with a mean of 2.28±0.30 (score 1-3), 2.16±0.40 (score 4-5) and 2.10±0.42 (score 6-9), respectively **(Figure 1B**).

### Faecal microbiota profile according to COVID-19 location of recovery

As an indirect measure of COVID-19 severity grade, the COVID-19 patients were grouped according with the location of recovery. Of the 111 COVID-19 patients with a characterised faecal microbiota, 59 (53.2%) required ICU admission, 39 (35.1%) were hospitalized in ward and 13 (11.7%) in ambulatory (household isolation). The gut microbiome composition of all COVID-19 patients were compared using the non-metric multidimensional scaling tool (**Figure 2A**). The faecal microbiota community of COVID-19 patients recovering in ambulatory is more similar between them than with the microbiota from those recovering in ward and in the ICU (p<0.05, PERMANOVA). The comparison of the relative abundance at phylum level between the three groups unveils a consistent trend for an increase in the relative abundance of Proteobacteria from 3% in ambulatory patients to 12% and 14% in ward and ICU patients, respectively (**Figure 2B**). The Firmicutes/Bacteroidetes ratio decreases in COVID-19 patients from ambulatory-ward-ICU (0.68, 0.65, 0.58, respectively). Like observed for the WHO severity groups, the COVID-19 patients hospitalized in ICU tends to have lower alpha diversity (Shannon’s index) in comparison with ambulatory and in ward/hospitalized COVID-19 patients **(Figure 2C)**, as suggested by the lower mean and the first and third quartile values.

**Figure 2.**
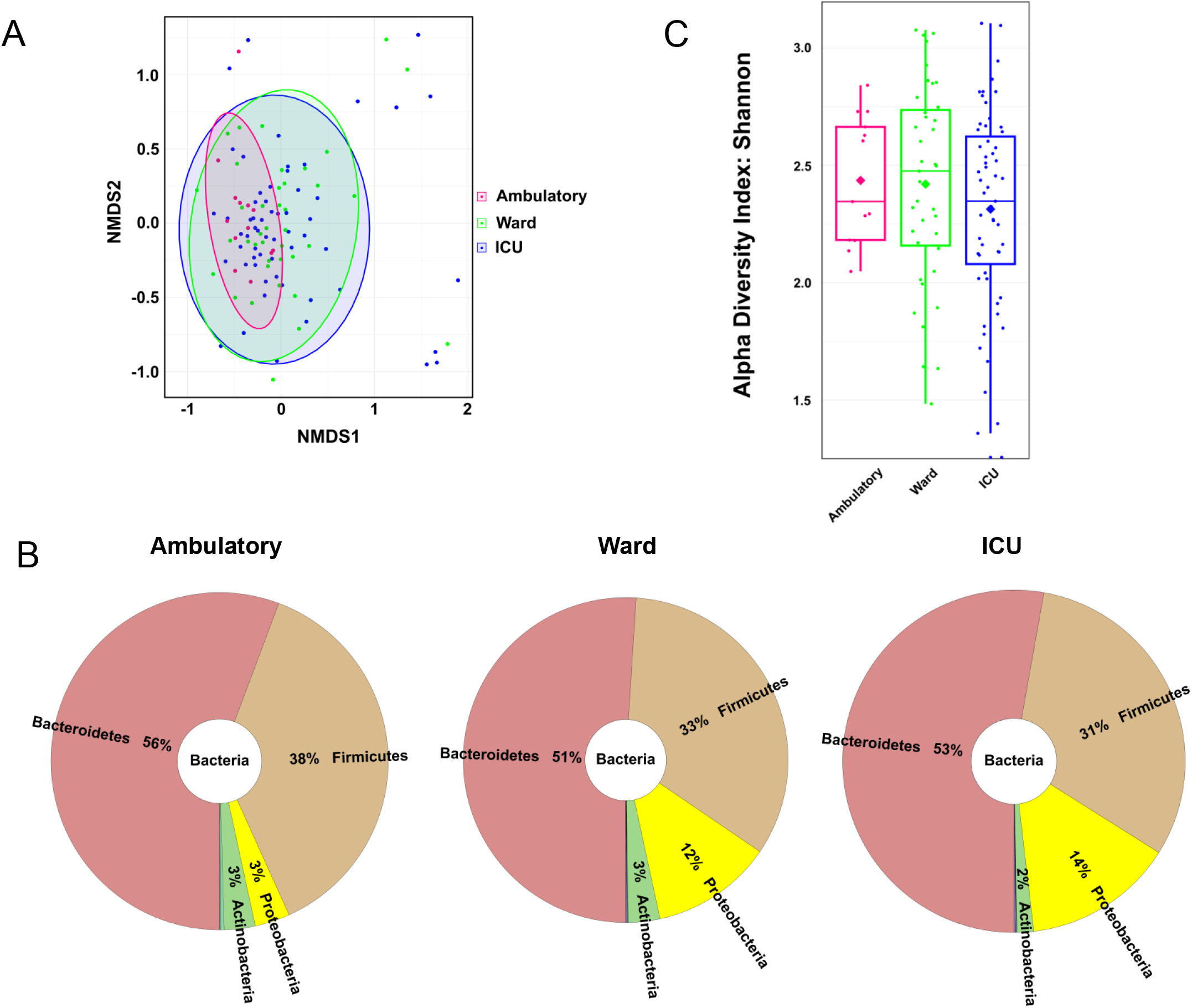
Faecal microbiota composition of COVID-19 patients according to patient location of recovery: ambulatory, hospitalized in ward or ICU. (A) Faecal microbiota community alterations according to patient location in NMDS2 (Non-metric multidimensional scaling) plot based upon Bray-Curtis dissimilarity. (B) Main bacterial phyla in faecal samples of COVID-19 patients according to patient location. (C) Boxplot of alpha-diversity (measured by Shannon’s diversity index) of COVID-19 patients according to patient location.

### Clinical characteristics associated with COVID-19 severity

Univariate and multivariate logistic regression models were used to evaluate associations between patients’ clinical characteristics and COVID-19 severity (**Table 2)**. Our aim was to develop a prognostic model able to predict the occurrence of certain outcomes in severely vs mild-to-moderately ill patients. The univariate model showed that severe COVID-19 patients were more likely to be men and to have elevated blood levels of CRP compared with mild-to-moderate COVID-19 patients. The association between men gender and higher severity of COVID-19 disease is observable by the higher proportion of men (72.9%) with severe COVID-19 disease in comparison with women. Age, body mass index, Shannon’s diversity index, comorbidities (hypertension and diabetes) and antibiotic therapy (at least once prior 6 months before COVID-19) were not significantly different between mild-to-moderate and severe patients. Regarding antibiotic therapy during the course of COVID-19, this variable was not significantly associated with COVID-19 severity (OR = 2.05; 95% CI [0.55-7.73]; p=0.287).

In the multivariate model that was mutually adjusted for CRP, Shannon’s diversity index, age and antibiotic therapy 6 months prior to COVID-19 diagnosis, the variables CRP and Shannon’s diversity index were significantly associated with COVID-19 severity while gender was no longer significantly associated (**Table 2**). Accordingly, the probability of having severe disease is 3.45 times higher when CRP levels ≥96.8 mg/L. Likewise, the probability of having severe COVID-19 symptoms is 2.85 times higher when Shannon’s diversity index is lower than 2.25. The geographic areas of the participating centres did not have impact on our multivariate regression model showing that disease severity and Shannon’s diversity index outcomes are centre-independent.

The discriminative/predictive power of the model was evaluated by the ROC-receiver operating characteristic-curve analysis. The receiver operating characteristics curve (ROC) analysis revealed an acceptable discriminative power of the model, with an area under the curve (AUC) of 0.707 (95% CI, 0.600–0.814) (**Figure S1**). Furthermore, our model correctly predicts 56.4% and 78.9% of patients with mild-to-moderate and severe disease, respectively.

### Faecal microbiota profile in patients positive for SARS-CoV-2 in faeces

Regarding that some authors suggest that faecal microbiota alterations are associated with the presence of SARS-CoV-2 in the gastrointestinal tract (18, 19, 24), we analyse for the presence of SARS-CoV-2 RNA in faeces. Sufficient amount of good quality faecal RNA to detect SARS-CoV-2 RNA in 112 patients (97.4%) among 115 recruited patients. From the 112 samples analysed, 45 tested positive (40% of the COVID-19 patients). Interestingly, the virus was detected mostly in men than in women (61.3% and 38.7% respectively, p<0.05). We then investigated if the presence of the virus in faeces was associated with changes in gut microbiota composition. As depicted in **Figures 3A**, no major differences were found in the distribution of the most abundant phyla and genera between patients positive and negative for SARS-CoV-2 in faeces.

**Figure 3.**
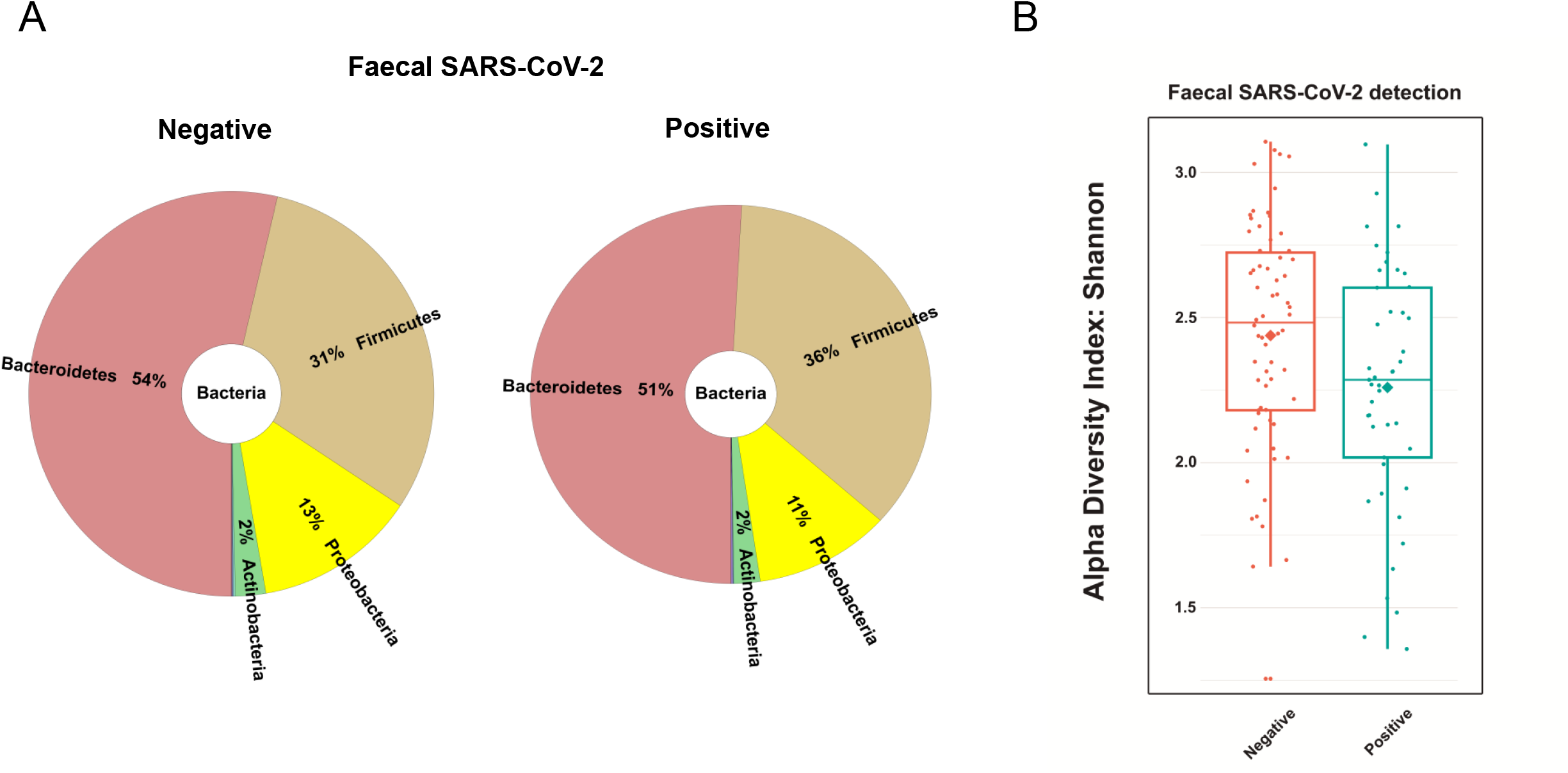
Faecal microbiota composition of COVID-19 patients according to the presence of SARS-CoV-2 in faecal samples. **(A)** Main bacterial phyla and **(B)** Boxplot of alpha-diversity (measured by Shannon’s diversity index) of COVID-19 patients according to the presence of SARS-CoV-2 in faecal samples.

**Figure 4.**
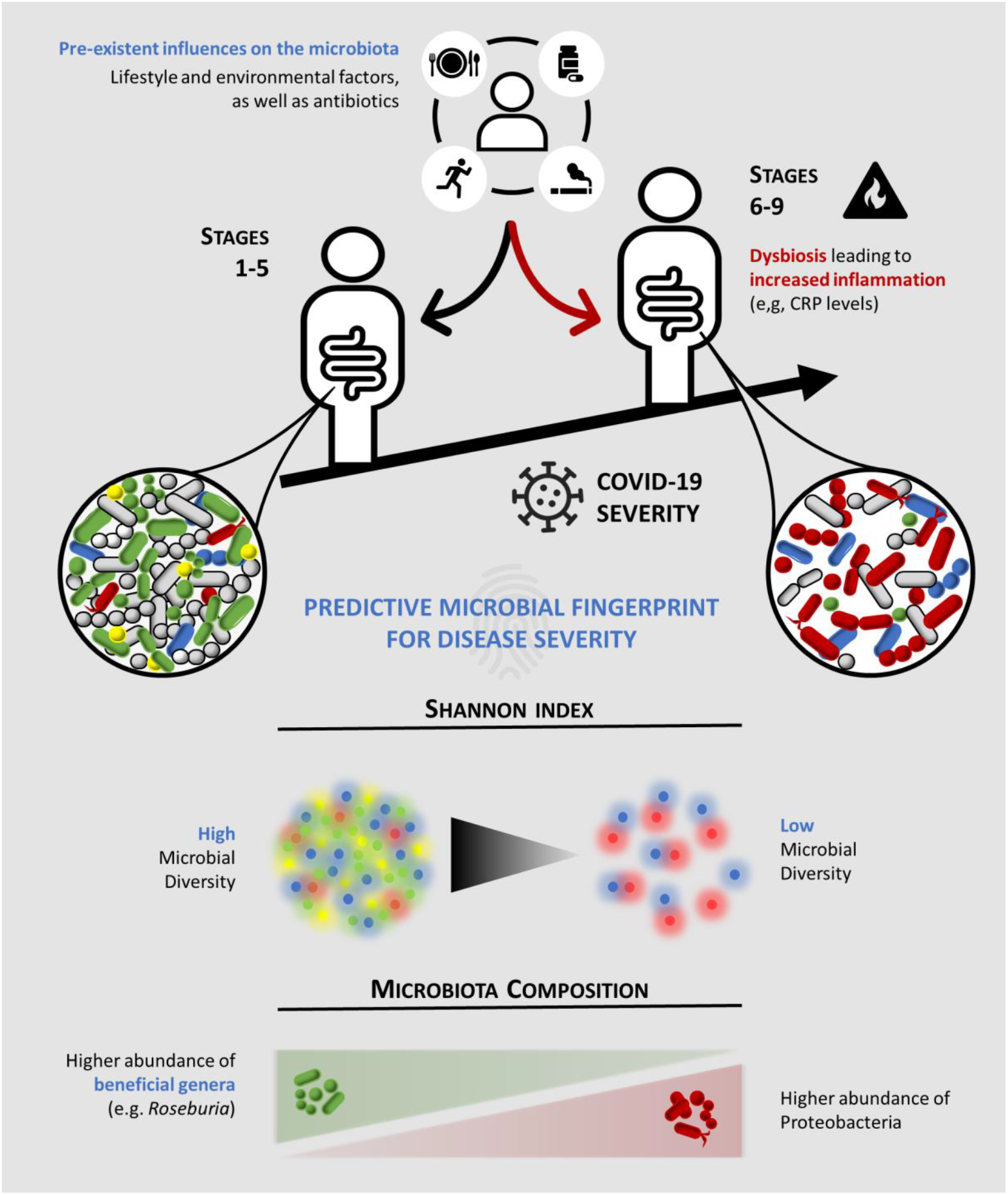
Schematic representation of the predictive microbial fingerprint for COVID-19 severity. Pre-existent influences on the microbiota, such as lifestyle and environmental factors, as well as antibiotics can induce dysbiosis (red arrow) leading to increased inflammation (e.g. CRP levels). Hence, a lower overall microbial diversity and abundance of beneficial commensal microorganisms (e.g. *Roseburia*), along with increased abundance of Proteobacteria are associated with severe COVID-19 severity (a score of ≥6 in WHO clinical progression scale). CRP, C-reactive protein.

Subsequently, we assessed the association between the faecal SARS-CoV-2 positivity and COVID-19 severity score or location of recovery using Pearson’s chi-square test. Importantly, no association was verified between the two categorical variables (p-value is 0.31 and 0.57 for severity score and location of recovery, respectively). Nevertheless, we found a strong tendency for a lower Shannon’s diversity index in faeces of SARS-CoV-2 positive patients (p=0.06) (**Figure 3B**).

## Discussion

We conducted a multicentre prospective cross-sectional study with 115 COVID-19 patients of different COVID-19 severity stages under the hypothesis that gut microbiota dysbiosis plays a pivotal role in the pathophysiology of COVID-19 namely in the severity of its clinical course.

In order to determine the association between the gut microbiota composition and COVID-19 disease severity, clinical and 16S rRNA gene sequencing data from COVID-19 patients were analysed and subsequently clustered according with: i) severity of COVID-19 using the WHO Clinical Progression Scale *i*.*e*. mild, moderate or severe; and ii) location of recovery from COVID-19 *i*.*e*. ambulatory, hospitalized in ward or ICU. Our data show for the first time an inverse association between relative bacterial abundance at genus level and Shannon’s index diversity with COVID-19 disease severity. According with our multivariable model, CRP ≥ 96.8 mg/L and Shannon’s diversity index <2.25 were associated with higher severity (a score of 6 or more in COVID-19 WHO clinical progression scale) suggesting that these patient’s variables are predictors for severe COVID-19. Indeed, our multivariable model correctly predicts 79% of patients with severe COVID-19.

Interestingly, faecal SARS-CoV-2 is detected in COVID-19 patients that tend to have lower Shannon’s diversity (p=0.06). We did not detect an association between the faecal SARS-CoV-2 positivity and COVID-19 severity score (p=0.31), however we cannot exclude that negative faecal SARS-CoV-2 patients could become positive during COVID-19 disease progression.

In comparison with mild COVID-19 patients, the gut microbiota from moderate and severe COVID-19 patients tend to have: **1)** decreased Firmicutes/Bacteroidetes ratio (0.68 in mild compared to 0.65 and 0.58 in moderate and severe COVID-19, respectively); **2)** higher abundance in Proteobacteria; (3% in mild compared to 12 and 14% in moderate and severe COVID-19, respectively); **3)** lower abundance of butyrate-producing bacteria from Lachnospiraceae family in particular *Roseburia* and *Lachnospira* genera; and **4)** lower abundance of Actinobacteria phylum namely Bifidobacteria and *Collinsella* genus. All these alterations are well-known microbial signatures of dysbiosis in gut microbiota (25-28).

Commensal bacteria play a fundamental role in the homeostasis of both immune and inflammatory functions of the gut (29). Anaerobic bacteria from Lachnospiraceae family such as *Roseburia* and *Lachnospira* genera produce butyrate, a short-chain fatty acid known to exert anti-inflammatory effects in the intestinal epithelium (30). Despite not being butyrate producers themselves, *Bifidobacterium* species are able to cross-feed butyrate-producing bacteria through the secretion of fermentation end-products such as acetate (31). This may constitute a potential mechanism by which *Bifidobacterium* species (32, 33) counteract intestinal viral infections. Another mechanism might be related with their capacity to decrease the production of pro-inflammatory cytokines (*e*.*g*. tumor necrosis factor-alpha and interferon-gamma) and increase the production of anti-inflammatory cytokines (*e*.*g*. interleukins 4 and 10) (34). Taking all this into consideration, we propose that changes in gut microbiota composition observed in severe COVID-19 patients may eventually act as a trigger to promote mucosal inflammation and increased gut permeability to proinflammatory molecules. Consequently, this may induce a state of systemic inflammation since these patients exhibited higher levels of blood CRP, a recently recognized prognostic factor for COVID-19 severity (35). Likewise, blood CRP concentrations ≥96.8 mg/L is associated with a score of 6 or more in COVID-19 WHO clinical progression scale in accordance with our multivariate model. The increase of Proteobacteria, a proposed signature of disease (36) particularly of epithelial dysfunction (37), in severe COVID-19 patients sustains our observation of a relation between dysbiosis microbiota and severity of COVID-19 disease.

Interestingly, the COVID-19 men patients seemed more prone to severe disease when compared with COVID-19 women (p=0.032). This gender discrepancy that has been described in other clinical trials (38), might be explained by a higher expression of ACE2 (39) in intestinal epithelial cells. This protein receptor is required for SARS-CoV-2 binding, invasion and persistence in host epithelial cells (40). Furthermore, COVID-19 patients that tested positive for the presence of SARS-CoV2 in faeces were mostly men (p<0.05) which reinforces the involvement of intestinal ACE2 in the severity of the course of the disease.

Our findings are consistent with two previous cross-sectional studies with COVID-19 patients carried on Hong Kong (China) (18, 19). The similarity of our results, collected in Portugal (a south-western European country), with the geographically far distant Chinese population led us to conclude that gut microbiota dysbiosis is a bona fide predictor of COVID-19 disease severity and the microbiome-based risk stratification should be considered for management of SARS-CoV-2 infection susceptibility, in parallel with worldwide-scale vaccination against COVID-19. Thus, our study open perspectives for the development of therapeutic interventions that aim to correct dysbiosis in severe COVID-19 patients. These include, dietary modifications, administration of butyrate-producing probiotics or prebiotics and faecal microbiota transplantation from healthy donors (41), shown to be effective in recurrent *Clostridium difficile* infection (42). These interventions are expected to increase overall bacterial diversity and the abundance of commensal bacteria, thereby contributing to inhibit the overgrowth of bacteria from Proteobacteria phylum.

In summary, we revealed for the first time an association between the gut microbiota and WHO Clinical Progression Scale, which reflects patient trajectory during COVID-19 disease. Our data show that gut microbiota dysbiosis is present in moderate and severe COVID-19 patients in comparison with asymptomatic/mild patients. Importantly, the evidence from this study suggests that CRP and gut microbiota diversity are prognostic biomarkers for severe COVID-19. Notwithstanding, a cross-sectorial study including a larger population size is necessary to produce a more powerful multivariable logistic regression model.

## Acknowledgments

The study was promoted by the NOVA Medical School of Universidade NOVA de Lisboa, CINTESIS, and sponsored by the Fundação para a Ciência e a Tecnologia (FCT – project number 268_596883842) and BIOCODEX. The funders had no role in study design, data collection, data analysis, data interpretation, or manuscript writing.

## Funding

This work was supported by the Fundação para a Ciência e Tecnologia under Grant n°268_596883842 and BIOCODEX.

## Declaration of interests

All authors declare no competing interests.

## Supplementary file captions

**Figure S1**. Receiver operating characteristics curve (ROC) analysis.

## Notes

### Competing Interest Statement

The authors have declared no competing interest.

