## Supplementary figures and images for "Gut microbiota diversity and C-Reactive Protein are predictors of disease severity in COVID-19 patients"

### Supplementary Figure S1

Supplementary Figure S1

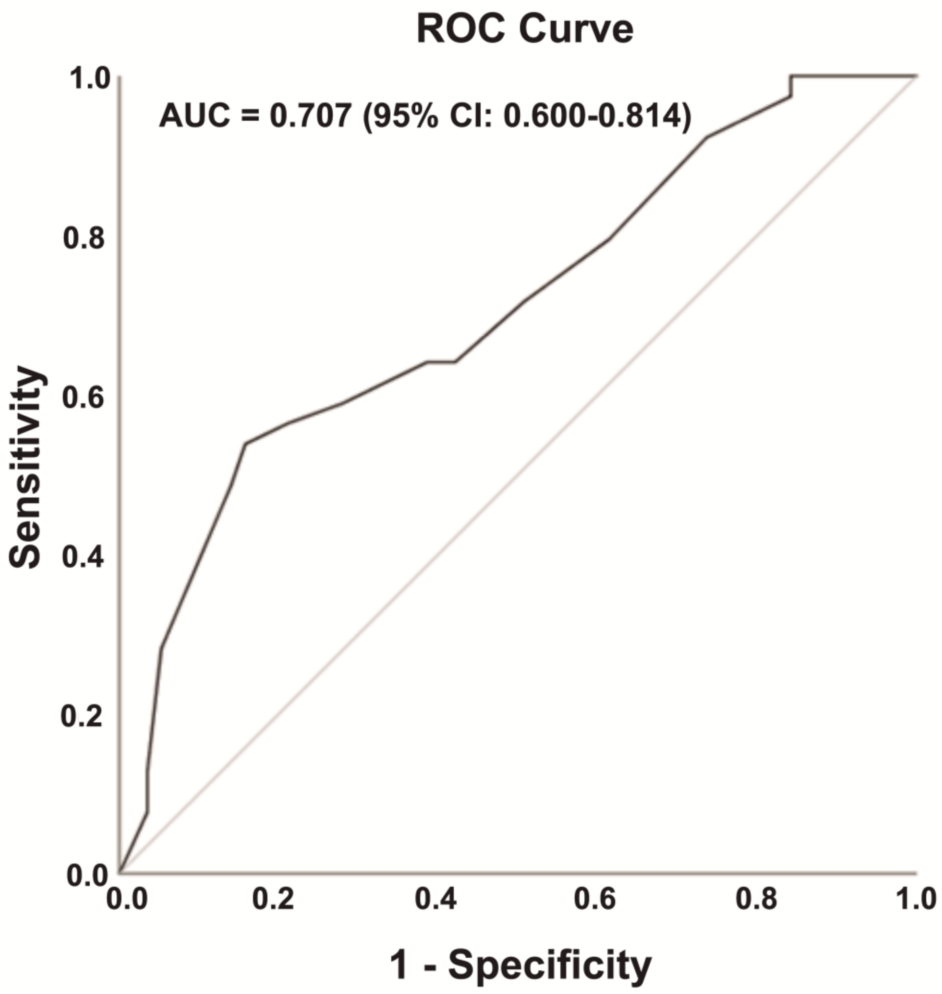
